# Dynamic Distribution Decomposition for Single-Cell Snapshot Time Series Identifies Subpopulations and Trajectories during iPSC Reprogramming

**DOI:** 10.1101/367789

**Authors:** Jake P. Taylor-King, Asbjørn N. Riseth, Manfred Claassen

## Abstract

Recent high-dimensional single-cell technologies such as mass cytometry are enabling time series experiments to monitor the temporal evolution of cell state distributions and to identify dynamically important cell states, such as fate decision states in differentiation. However, these technologies are destructive, and require analysis approaches that temporally map between cell state distributions across time points. Current approaches to approximate the single-cell time series as a dynamical system suffer from too restrictive assumptions about the type of kinetics, or link together pairs of sequential measurements in a discontinuous fashion.

We propose Dynamic Distribution Decomposition (DDD), an operator approximation approach to infer a continuous distribution map between time points. On the basis of single-cell snapshot time series data, DDD approximates the continuous time Perron-Frobenius operator by means of a finite set of basis functions. This procedure can be interpreted as a continuous time Markov chain over a continuum of states. By only assuming a memoryless Markov (autonomous) process, the types of dynamics represented are more general than those represented by other common models, e.g., chemical reaction networks, stochastic differential equations. Additionally, the continuity assumption ensures that the same dynamical system maps between all time points, not arbitrarily changing at each time point. We demonstrate the ability of DDD to reconstruct dynamically important cell states and their transitions both on synthetic data, as well as on mass cytometry time series of iPSC reprogramming of a fibroblast system. We use DDD to find previously identified subpopulations of cells and to visualize differentiation trajectories.

Dynamic Distribution Decomposition allows interpreting high-dimensional snapshot time series data as a low-dimensional Markov process, thereby enabling an interpretable dynamics analysis for a variety of biological processes by means of identifying their dynamically important cell states.

**Author summary:** High-dimensional single-cell snapshot measurements are now increasingly utilized to study dynamic processes. Such measurements enable us to evaluate cell population distributions and their evolution over time. However, it is not trivial to map these distribution across time and to identify dynamically important cell states, i.e. bottleneck regions of state space exhibiting a high degree of change. We present Dynamic Distribution Decomposition (DDD) achieving this task by encoding single-cell measurements as linear combination of basis function distributions and evolving these as a linear system. We demonstrate reconstruction of dynamically important states for synthetic data of a bifurcated diffusion process and mass cytometry data for iPSC reprogramming.

## Introduction

Data-driven reconstruction of dynamic processes constitutes a central aim of systems biology. High-dimensional single-cell molecularly resolved time series data is becoming a key data source for this task [1, 2]. However, these technologies are destructive, and consequently result in *snapshot time series* data originating from batches of cells collected at time points of interest. A longstanding and still challenging problem is to reconstruct dynamic biological processes from this data, to the end of identifying dynamically important states, i.e. regions of state space that cells preferentially pass through, these include transitionary states (bottlenecks) when differentiation decisions are made, and terminal states. With *snapshot time series*, it is challenging to identify these states as we cannot track the state of an individual cell from one time point to a new state at a later time point, one has to temporally map between state distributions.

Chemical reaction networks are a popular class of parametric models assuming that the temporal state evolution is well described by chemical kinetics. Ordinary differential equations (ODEs) are used to describe smooth deterministic dynamics, and stochastic differential equations (SDEs) for dynamics in the low copy number/concentration regimes affected by stochastic fluctuations. Chemical reaction network models require explicit definition of the model structure, i.e. set of reactions or interactions among the system components. This task is manageable for small, well defined systems, such as small signaling systems [3]. However, by means of high-dimensional measurements, we typically observe larger systems comprising at least dozens of components with largely *a priori* undefined interactions. This situation results in a combinatorial explosion of model variants that cannot be exhaustively evaluated [4, 5]. Alternative approaches are agnostic with regards to parametric form and model structure and use a probabilistically-motivated rule to map between distributions, e.g., one optimal transport method maps neighbours at one time point to the nearest neighbour at the next time point [6]. However, such generic approaches are rather extreme in their agnosticism and abandon reasonable assumptions on the dynamics of cellular systems, e.g., that cells can be modelled as an *autonomous* dynamical systems in continuous time, such as a Markov chain, where the cell’s current state infers its likely future state independent of the current time within the experiment.

Operator approximation methods constitute an alternative class of models that are agnostic to model structure and yet allow for encoding of general system properties such as autonomy, conservation of mass, and boundary conditions. These methods approximate both the Perron–Frobenius operator [7] and the Koopman operator [8]. These operators describe the evolution of distributions and other functions of a dynamical system’s state. The early theory on these operators was developed to describe systems in classical, statistical, and quantum mechanics [9–13], and in probability theory [14, 15]. The operators fully describe a nonlinear dynamical system as a linear system in higher, possibly infinite, dimensions. Hence, techniques from linear analysis can be utilised to gain insight from the systems; in particular, calculation of eigenvalues and eigenfunctions allow for timescale separation. Eigenfunctions of linear operators show the fundamental building blocks of possible behaviours available to a dynamical system, e.g., exponential growth/decay, oscillations, a steady state. Data-driven approximations to the operators have been investigated in the past years, originating in the computational fluid dynamics community [16–20]. Their focus is to approximate finite-dimensional projections of the Koopman operator with a family of algorithms known as Dynamic Mode Decomposition (DMD). The algorithm has further been applied to other areas such as neuroscience, infectious disease epidemiology, and control theory [21–23] and parameter estimation [24, 25]. When carrying out approximations of these operators, eigenvalues can be ordered in terms of magnitude to extract slow behaviour (approximated well) to fast behaviour (representative of noise) [25]. Dynamic Mode Decomposition assumes that the data is recorded at equally spaced time points, whilst our work extends the technique to support data recorded at arbitrary time points.

We adapt Dynamic Mode Decomposition to identify dynamically important states from single-cell snapshot time series. Our method is based on representing the distribution at each time point via basis functions and calculating an approximation to the Perron–Frobenius operator by minimising an error term — akin to least squares when fitting ODEs. The error terms then give an indication of how well the data fits into the model assumptions: primarily that the data is generated by an autonomous dynamical system. Because the Perron–Frobenius operator describes the evolution of distributions, we name our approach Dynamic Distribution Decomposition (DDD) in keeping with the DMD naming convention. DDD leads to the calculation of a Markov rate matrix but over a continuum of states — as opposed to discrete states. As previously mentioned, one can then use standard methods of analysis for linear operators based on eigen-decompositions. As Markov processes can be represented as directed weighted graphs, our graph can be evaluated in two dimensions and then the high-dimensional operator and its corresponding eigenfunctions have a natural low dimensional representation. Our approach also allows for visualisation of inferred state trajectories as a branching structure when cell fates are stochastic, and approximation of fitting error when matching model prediction to sample data. By using a Markov rate approach over distributions, we overcome the difficulties listed above. We demonstrate DDD on a synthetic stochastic dynamical system representing cells making a cell differentiation decision as well as for a mass cytometry time series taken from an iPSC reprogramming of a fibroblast cell line taken from Zunder *et. al*. [26] to re-identify subpopulations of cells as first elucidated in the original manuscript.

## Results

### Inference of State Distribution Dynamics by Approximation of the Perron– Frobenius Operator

We developed a method herein referred to as Dynamic Distribution Decomposition to analyse snapshot time series, consisting of the following stages (illustrated in Figure 1): (*a*.) data at each recorded time point *t*_1_, …, *t_R_* are fitted to a set of basis functions and (*b*.) encoded into coefficient vectors *c*_1_, …, *c_R_*; (*c*.) a fitting procedure is carried out to infer the most likely continuous linear map between the coefficients, generating fitting errors *ε*_1_, …, *ε_R_*; (*d*.) eigenfunctions are then analysed; and (*e*.) in high dimensions graph based visualisations can be used for eigenfunctions, for full details see Methods section. In the case where probability density functions are used as basis functions, the linear map denoted *P* can be interpreted as a Markov rate matrix; the structure of this matrix is often dense but its dominating structure can be elucidated via Lasso regularisation, see Figure 1(f). We applied our method to two systems: first, simulated particles in a potential well; and second, experimental data of iPSC reprogramming of a fibroblast system.

**Fig 1.**
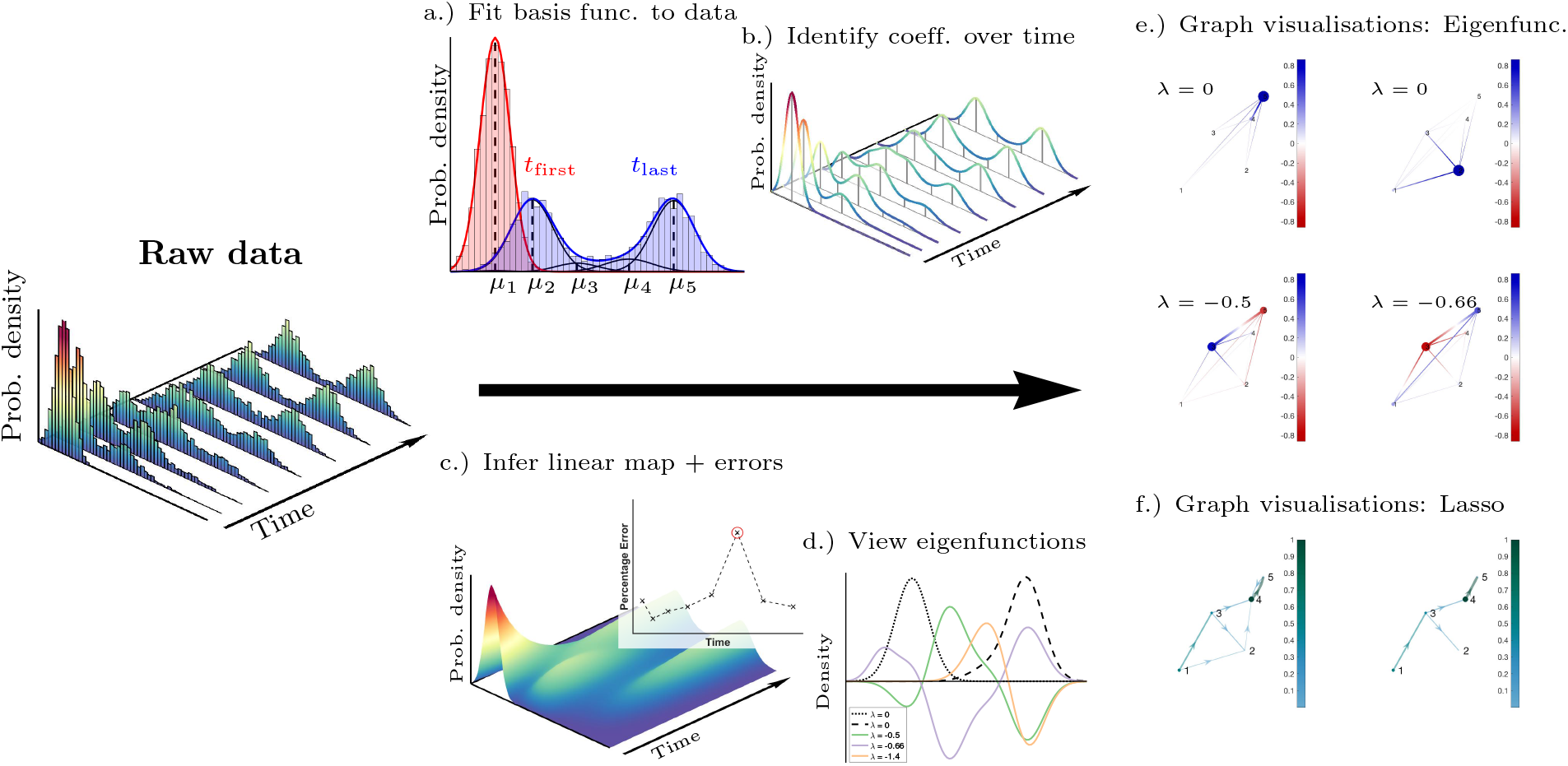
Illustration of DDD workflow. (a.) Gaussian mixture models as basis functions: single-cell profiles from each time point are fitted to a basis of Gaussian mixture model, two distributions shown in red and blue; (b.) identification of the coefficients *c*(*t*) at each time point, *t* = *t*_1_, …, *t_R_*; (c.) the Perron–Frobenius matrix *P* enables us us to infer the likely state for all time points and generate errors *ε*_1_, …, *ε_R_*; (d.) examination of the eigenfunctions in low dimensions; or in high dimensions (e.) using graphical visualisations; and (f.) a Lasso regularisation can reveal sparse structure.

### Particles in Potential Well with Fluctuations

The first numerical example is for illustrative purposes whereby we know the stochastic process generating the sample points. We consider simulated particles in a bistable potential well undergoing fluctuations. After initialisation around point (1, 1)^τ^/2, particles stochastically switch between one of two paths: *y* = 2*x* or *y* = *x*/2 to finally settle in one of the two final state (2, 4)^τ^ or (4, 2)^τ^. We model this process by the two-dimensional SDE 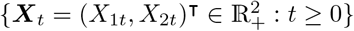 

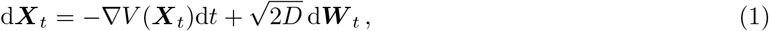

where ***W***_t_ is a two-dimensional Wiener process. The potential well is of the form 

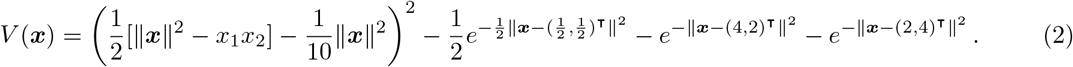

As an initial condition at *t* = *t*_1_, the sample is placed with a multivariate normal distribution with mean *µ* = (1, 1)^τ^/2 and covariance matrix = Σ *I*_2_/2 where *I*_2_ is the identity matrix. The diffusion constant is chosen to be *D* = 1/4. Along the lines *x* = 0 and *y* = 0, the system has reflecting boundaries imposed. For simulations, the Euler–Maruyama numerical scheme is used^1^ with time step *δt* = 2^−9^. Three sample trajectories are visualised, see Figure 2(a); and the potential well is plotted, see Figure 2(b).

**Fig 2.**
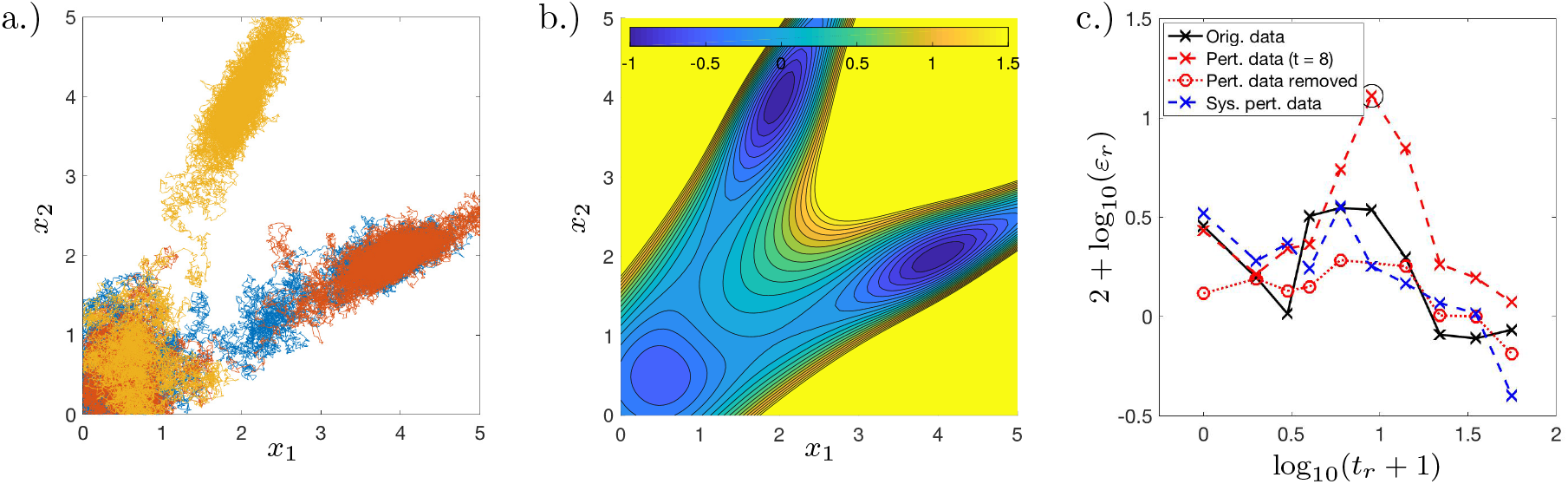
(a.) Three simulated trajectories are shown from the stochastic dynamical system described by equation (1)–(2). (b.) The potential well as described in equation (2) is shown. (c.) Log-log error plot showing time point against log_10_ of the percentage error; the original data set(black solid line) has a mean error of 2.0%; the perturbed data set (dashed red line) has a mean error of 3.9%; the perturbed data set with the erroneous data point removed (dotted red line) has an error of 1.3%; and the data set with systematic error (dashed blue line) has a mean error of 1.9%.

The system is simulated 2000 times and observed at the time points *t* = 0, 1, 2, 3, 5, 8, 13, 21, 34, 55. The time points are only partially observed where half the samples are (uniformly) randomly discarded. Using Gaussian mixture models of various sizes, we choose to use 3 basis functions for each time point totalling *N* = 30 basis functions.

### Extrema of Eigenfunctions Identify Steady State and Bistable Paths

The eigenfunctions of the approximated Perron–Frobenius operator allow us to identify the steady state and bistable paths in the above system. In low dimensions we visualise eigenfunctions of *P* as a continuous function, see Figure 3(a,b,c); or a graph, see Figure 3(d,e,f) and Methods section. Eigenfunctions corresponding to eigenvalues with large absolute value are approximated with larger error than in cases with small eigenvalues, a point also noted in Ref. [25]; notice in Figure 3(c,f) there are a few small fluctuations around (1, 1)^τ^/2. Also, eigenvalues and eigenfunctions are basis function dependent, so changes in basis functions change the eigen-decomposition. However, regardless of changes to the basis functions, the key dynamic states (as visible in the eigenfunctions) remain the same provided the changes to the basis functions are not drastic. Since we use 30 basis functions, hypothetically we can find 30 eigenfunctions. However, we just plot the first three eigenfunctions; these are real with no imaginary component. From these figures, it is clear that three basins around (2, 4)^τ^ and (4, 2)^τ^ and at the initial condition (1, 1)^τ^/2 are dynamically important. Therefore, examination of the first few eigenfunctions allows for detection of dynamically important states.

**Fig 3.**
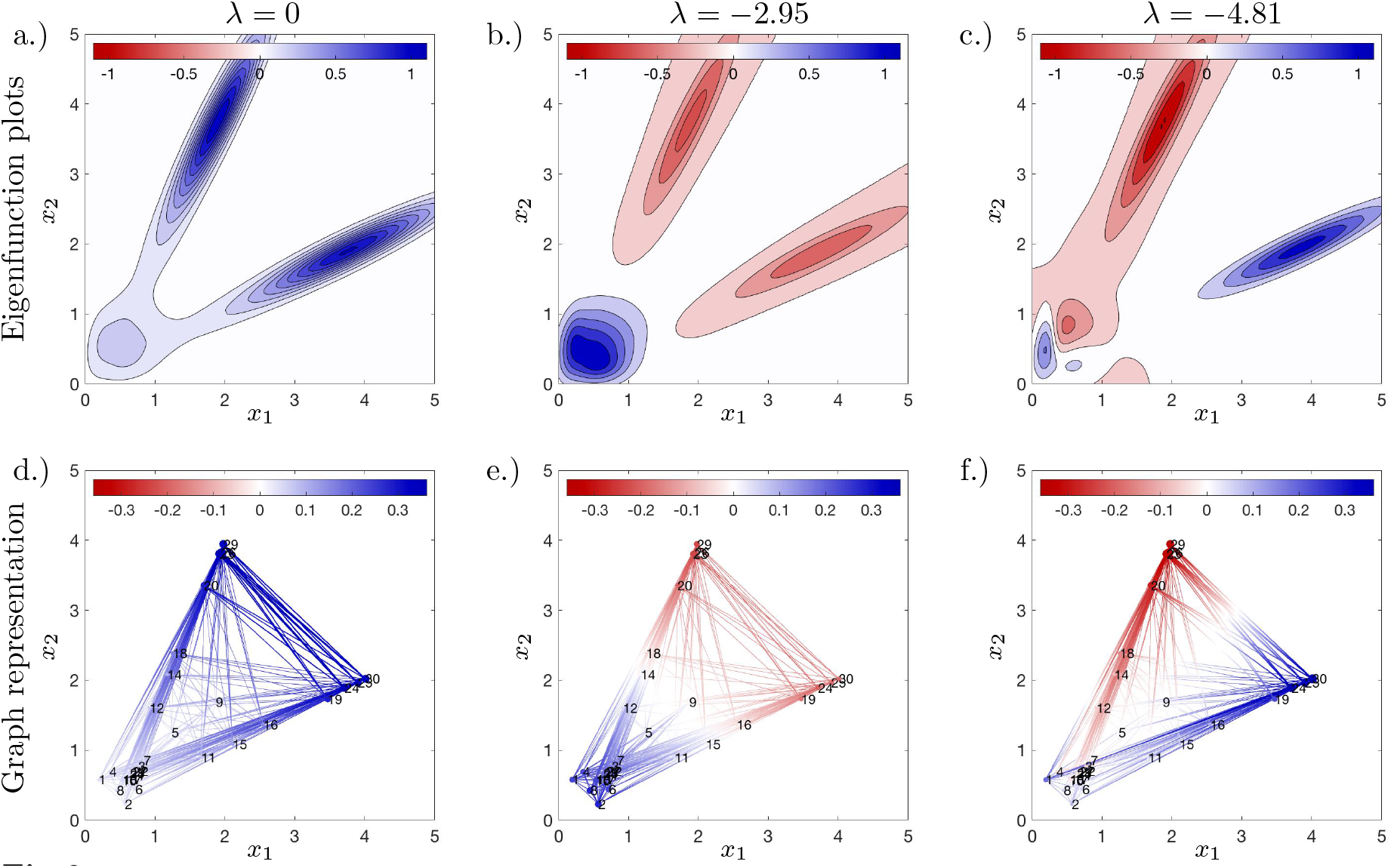
Dynamic distribution decomposition applied to data generated by stochastic dynamical system described by equation (1)–(2). Plots (a)–(c) show a plot of the two dimensional eigenfunction, and plots (d)–(f) shows a corresponding graph representation of the eigenfunctions, for full details see Methods section.

### Dynamic Distribution Decomposition is Robust to Noisy Observations

We evaluated the robustness of our inference procedure to measurement noise. Specifically, three further modified data sets are also considered: (*i*.) considering a single time point perturbed by additive random noise; (*ii*.) removing this perturbed time point and fitting the model; and (*iii*.) randomly perturbing all time points by additive random noise. For all perturbations, sample points are modified by an additive error term drawn from a zero-mean multivariate Gaussian with covariance matrix Σ = *I*_2_/4.

We plot the time points transformed by log_10_(*x* + 1) against the log-percentage error, i.e., 2 + log_10_(*ε_r_*) for *r* = 1, …, *R*, see Figure 2(c). We find that all data sets have consistently low error, but with an increase in error at the beginning of the realisation; this is due to the boundary conditions which were not incorporated into the choice in basis functions, which one would typically do when solving PDEs via a Galerkin approximation. The data set perturbed at time *t* = 8 (dashed red line) leads to increased error immediately before and after this time point (circled in black); after removing this erroneous data (fitting without *t* = 8) one obtains reduced errors comparable to the original data set (dotted red line). When adding systematic error to all time points (dashed blue line), one observes similar errors to the original data set; the reason for this is that the Gaussian basis functions now have a covariance matrix with larger entries (whilst the means remain similar). The eigenfunction plots are also similar to those generated by the original data set but more spread out (not plotted). Therefore we see that DDD is robust with regards to noisy observations.

### Lasso Regularisation Reveals Sparse Topology

We utilize Lasso regularization to identify key transition states, see Methods section. Specifically, *P* as a Markov rate matrix with the nodes located at the mean of the components of the Gaussian mixture model. The resulting network is cluttered and is hard to identify meaningful states or transition, see Figure 4(a). Lasso regularisation encourages sparsity and reveals the simple underlying structure, see Figure 4(b). The skeletal structure shows that around the initial condition the particle becomes strongly committed to one branch over the other — an accurate reflection of the dynamical system.

**Fig 4.**
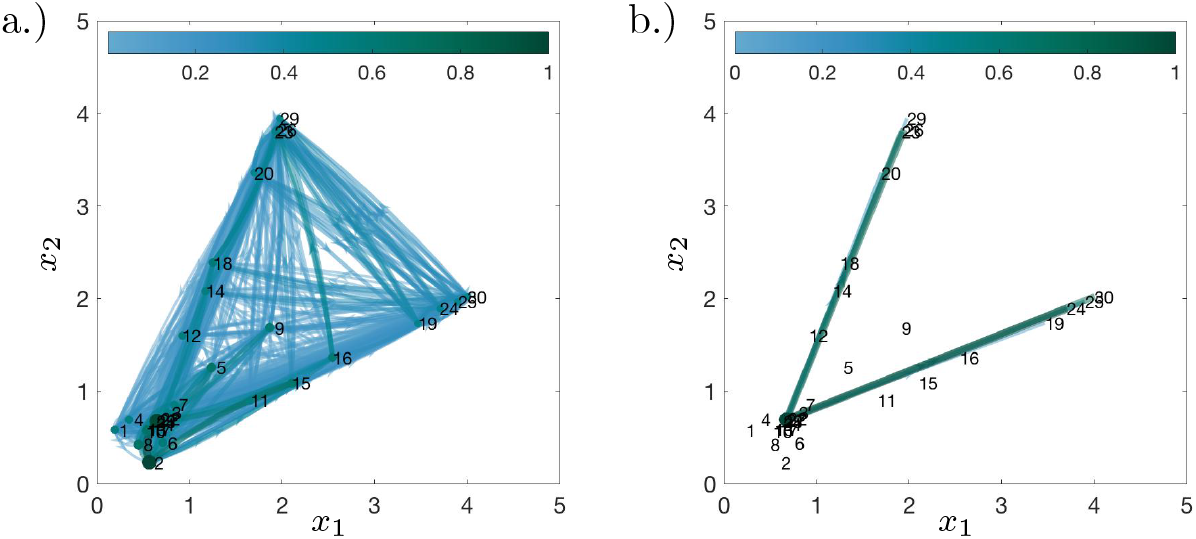
Dynamic distribution decomposition applied to data generated by stochastic dynamical system described by equation (1)–(2). The representation of *P* as a Markov rate matrix, colours of edges denote magnitude of rate change from one node to another, and node colours denote rate at which node decays. Colours scaled to unit interval. (a.) Markov transition matrix *P* plotted without Lasso regularisation; and (b.) Markov transition matrix *P* plotted with Lasso regularisation, β = 1/[100 × mean(*M*)], see Methods section.

## Mass Cytometry Data: iPSC Fibroblast Reprogramming

We studied the process of iPSC reprogramming using Dynamic Distribution Decomposition. We considered data from a study established by Zunder *et. al*. [26]. Specifically, the reprogramming of a fibroblast cell line differentiating into an induced pluripotent stem cell state was studied using mass cytometry. Cells are labelled using mass-tag cell barcoding, stained with antibodies before being measured via CyTOF. We focus our study to the cell line with the largest amount of cell events, i.e. on a Nanog-Neo secondary mouse embroyic fibroblasts (MEF) that expresses neomycin resistance gene from the endogenous Nanog locus. Reprogramming was monitored by Dox induction for 16 days followed by subsequent addition of LIF; the experiment was carried out over 30 days. Experiments were initialised together and cells harvested every 2 days until 24 days with a final measurement taken at the final time point; 18 protein markers were used as proxies to measure pluripotency, differentiation, cell cycle status, and cellular signalling.

We now briefly state our method for choosing basis functions. In the synthetic data example, we used prior information that there were 3 clusters, so used a 3 component Gaussian mixture model for each time (therefore *N* = 30 for 10 time points). For non-synthetic data we do not necessarily have this information, therefore we developed an approach to choose an expressive set of basis functions without letting their number grow too large and thereby ensure efficient solving of the minimisation problem later presented in equation (17). We fit multiple Gaussian mixture models to each time point, varying the number of components until the AIC curve flattened out [27]; in our case this happens at approximately 8 basis functions per time point. To avoid overfitting, we use regularisation to specify minimum diagonal entries of the covariance matrix. As our data has been scaled via the commonly used transformation function *f*(*x*) = arcsin(*x*/5) and then standardised by *z*-scoring, we choose a regularisation value of 1/2; smaller values can be used should one wish to capture sharp peaks, but at the cost of additional basis functions. Finally, for each time point we cluster the data into these Gaussian mixture models and remove poorly populated components. Here, after clustering, we remove any basis functions which represent less than α × 100% of the data. Therefore, it should be noted that obtaining a good fit to the matrix *P* is a payoff between: (*i*.) number of basis functions per time point (i.e., what is the maximum number of clusters per time point?); (*ii*.) the regularisation value (i.e., how sharp peaks can one fit?); and (*iii*.) the drop rate α (i.e., what fraction of data points does each basis function have to represent?).

We now decrease α and evaluate whether we have sufficient basis functions. We plot the percentage fitting error at each time point and the mean percentage error as a function of α, see Figure 5(a,b). These figures show that as α decreases, the error only minimally decreases for large increases in the total number of basis functions *N*. We can also view the eigenvalues plotted in the complex plane for various values of α, see Figure 5(c). We rescaled time to the unit interval, therefore one will not be able observe eigenfunctions with a corresponding non-zero eigenvalue ℛ(λ) > −1, i.e., we cannot observe timescales slower than the observation window. We notice for α = 0.005 that ℛ(λ_1_) = −1.06 so we are confident decreasing α will not offer much benefit. Additionally in the cases where α = 0.01 and α = 0.005, the extrema of the first few eigenfunctions correspond to the same basis functions not plotted.

**Fig 5.**
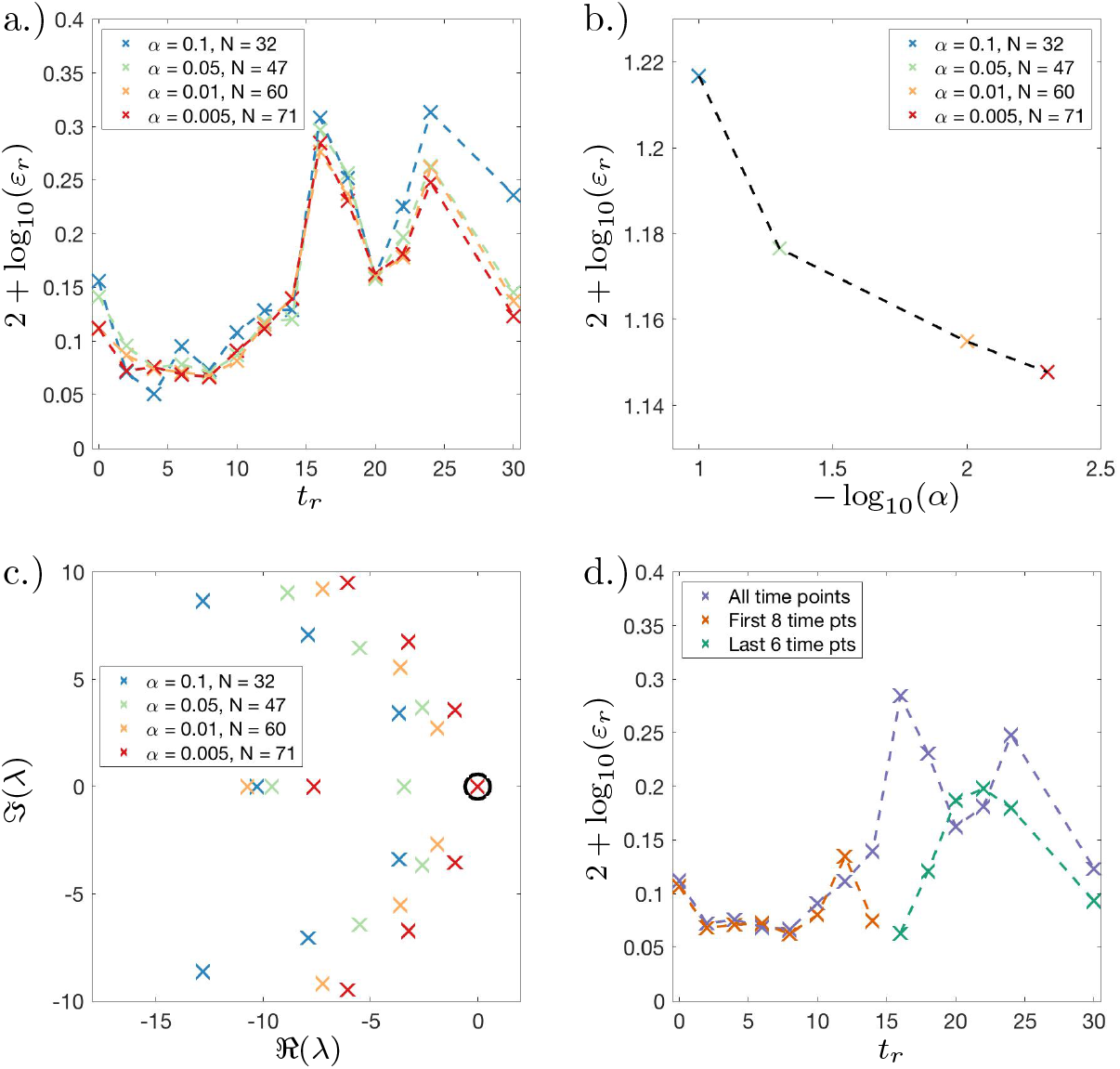
Dynamic distribution decomposition applied to Nanog-Neo cell line data taken from Zunder *et. al*. [26]. (a.) Fitting error plot showing log_10_ transformed percentage error plotted against time for different values of α (described in main text); (b.) log-log plot of mean fitting error plotted against −log_10_(α), mean error decreases as alpha decreases; (c.) complex plot of eigenvalues λ of Perron–Frobenius (PF) matrix *P* for different values of α; (d.) Fitting error plot showing log_10_ transformed percentage error plotted against time for α = 0.05, two further PF matrices are fitted using only the: first 8 time points; and last 6 time points.

### Loss of Dynamic Autonomy after Stimulus Removal

When a stochastic dynamical system is autonomous, the current state of the system determines the likely future states; here we show that after and including *t* = 16 days the system becomes less autonomous, once the Dox induction had ended. We see the error plotted again at each time point for α = 0.005, see Figure 5(d). We notice that time points after and including *t* = 16 days contain the vast majority of fitting error. To rule out the possibility that the dynamical system instantaneously changed at *t* = 16 days, we fit two Perron–Frobenius matrices, one using the first 8 time points and a second using the last 6 time points with all fits using the same basis functions. We find that there is still much more error contained in the final 6 time points compared to the first 8.

The autonomous dynamical system assumption means that using the data presented, the future states of the system depend on the current state. While this is likely true within a cell culture system, we only observe a tiny fraction of the state space of the dynamical system as we do not measure the transcriptome and the vast majority of the proteome. Therefore, it seems reasonable to assume that from *t* = 16 days, we are not observing enough of the dynamical system to obtain a linear map between distributions. This insight suggests further single-cell experiments at these later time points using technologies allowing greater ‘omic’ profiling, e.g., single-cell RNA-Seq.

### Inferred Dynamically Important States Agree with Previously Described Cell Subpopulations

We evaluated the extreme of the eigenfunctions of the approximated Perron–Frobenius operator to re-identify cell subpopulations found in Zunder *et. al*. [26]. We first plot the first 6 eigenfunctions, see Figure 6. Nodes that are close together to each other in 18 dimensions (using Euclidean distance) as plotted as close to each other in 2 dimensions. Protein expression of the basis functionsare also plotted using the same coordinates as the graph, see extra figure in Appendix.

**Fig 6.**
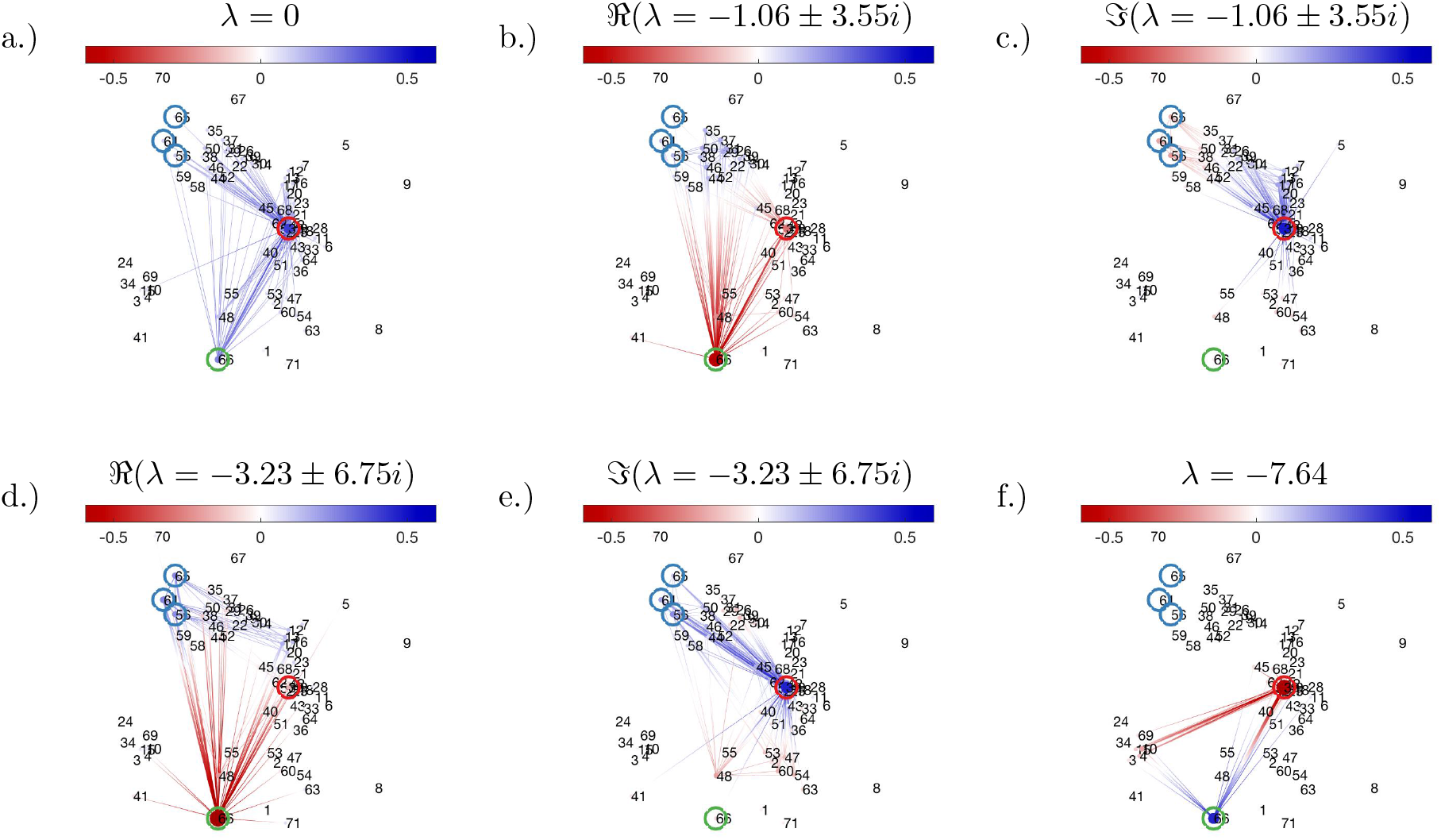
Dynamic distribution decomposition applied to Nanog-Neo cell line data taken from Zunder *et. al*. [26]. Groups as defined in the text are circled and coloured in: red (group A, MEF-like); blue (group B, mesendoderm); and green (group C, ESC-like).

When examining the extrema of the eigenfunctions, basis functions seem to cluster in 3 groups: group A centred around basis function 32; group B with members 56, 61, and 65; and group C with one member, basis function 66. Our algorithm recovers the same populations as stated in Zunder *et. al*. [26]: cells with low Ki-67 expression do not successfully reprogram and remain MEF-like (group A); cells with high Ki-67 expression then subdivide into two populations, an embryonic stem cell-like (ESC-like) population with Nanog^+^, Sox2^+^, and CD54^+^(group C) and a mesendoderm like population with Nanog^−^, Sox2^−^, Lin-28^+^, CD24^+^expression (group B). As our basis functions were added sequentially per time point, the MEF-like population appeared first.

DDD suggests a few new insights previously not elucidated in Zunder *et. al*. [26]. We find according to the fitted Perron–Frobenius operator, MEF-like cells form the steady state (when λ = 0). Therefore, the model predicts all cells would revert to fibroblasts if enough time passes — although one has to be careful over interpreting predictions due to the higher error after *t* = 16 days.

### Lasso Regularisation Reveals Sparse Topology of iPSC Dynamics

Reminiscent of the SDE example, the graph as induced by the transition matrix *P* is cluttered due to an abundance of low weighted edges, see Figure 7(a); we apply the Lasso modification to reveal a two branching points, see Figure 7(b) and Methods section. Finally, to focus on the 3 groups previously identified, we prune edges leading to unannotated nodes to obtain an easily interpretable branching structure, see Figure Figure 7(c). This figure suggests that at basis function 53 (close to basis function 1, i.e., the initial state), a cell moves to towards branching basis function 16 (CD73^−^, CD140a^+^, CD54^+^, Oct-4^+^), and then has a decision to move towards basis function 32 (group A, MEF-like) or to reach a second branching point at basis function 29 (CD73^+^, CD140a^+^, CD54^−^, Oct-4^+^, KLF4^+^). At basis function 29, the cell will then choose between basis functions 56, 61 and 65 (group B, mesendoderm population), or towards basis function 66 (group C, ESC-like); there is also a weakly weighted edge back to basis function 32 (group A, MEF-like). The state described by basis function 29 was previously described in Zunder *et. al*. [26], but we are able to include an additional transitionary state by means of basis function 16. We can conclude that the cell decision towards becoming remaining MEF-like is made early during the course of the experiment: basis function 16 was placed with the data recorded at 6 days; basis function 29 was placed with the data recorded at 12 days.

**Fig 7.**
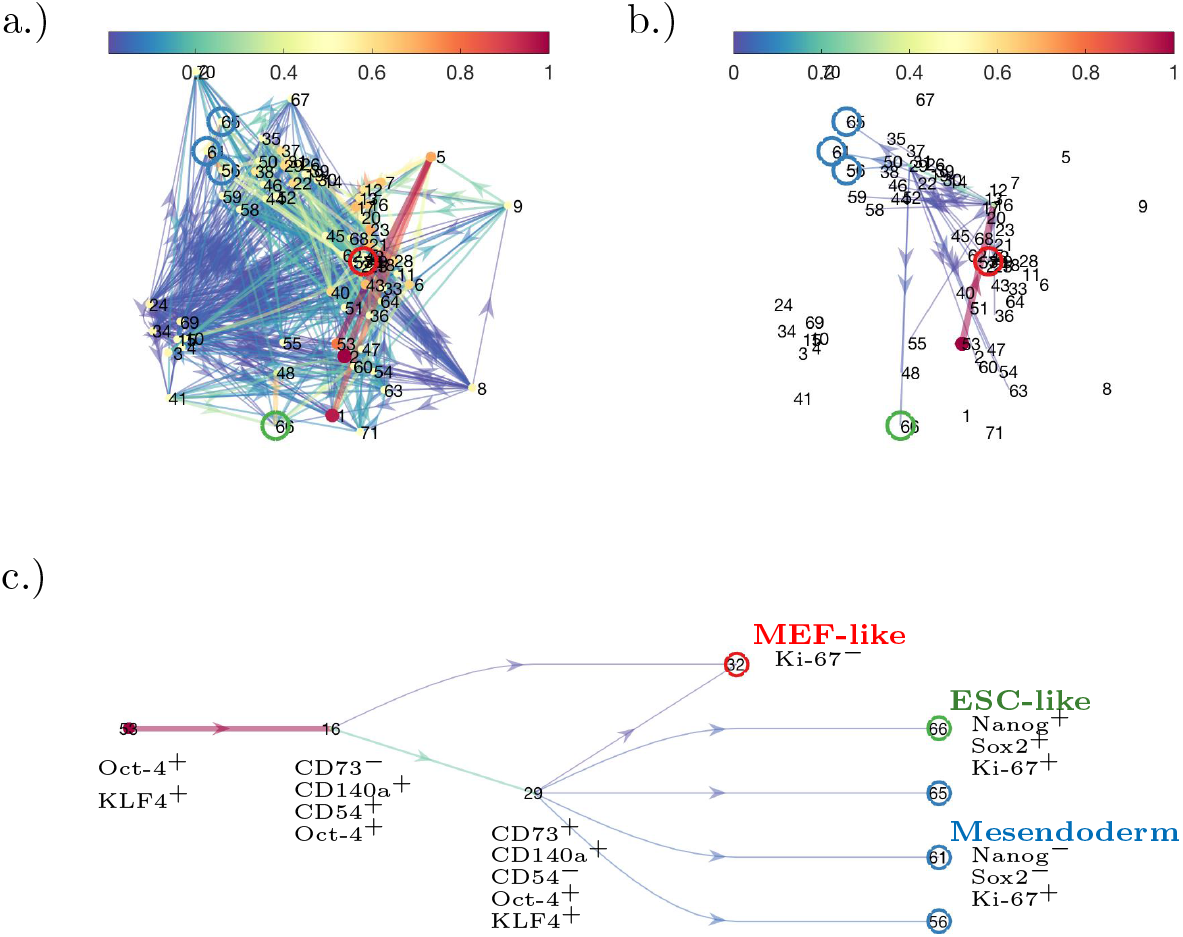
Dynamic distribution decomposition applied to Nanog-Neo cell line data taken from Zunder *et. al*. [26]. (a.) Markov transition matrix *P* plotted without Lasso regularisation; and (b.) Markov transition matrix *P* plotted with Lasso regularisation, β = 1/[800 × mean(*M*)], see Methods section. In (c.) the graph structure from (b.) has unannotated end nodes removed and is rearranged into a simple branching structure. Groups as defined in the text are circled and coloured in: red (group A, MEF-like); blue (group B, mesendoderm); and green (group C, ESC-like).

## Methods

We now give the mathematical set-up to our problem, additional technical details are given in the appendix. The method follows the following steps: (*i*.) the statement is posed that the temporal evolution of cell states follows a linear partial differential equation; (*ii*.) the distribution of the sample points at each time point can be encoded into a sequence of basis functions; (*iii*.) the weights of these basis functions can change dynamically interpolating between sample points; and (*iv*.) we fit the form of the matrix approximation of differential operator around these changing basis functions; and (*v*.) study the eigenfunctions. The workflow of the method is also as an illustration in Figure 1.

### Mathematical Set-Up

Assume we have a sequence of *R* experimental readings at time points *t*_1_ < … < *t_R_*; without loss of generality we choose *t*_1_ = 0. At each time point, *n_r_* cells are harvested with states *X_r_* = {*x_r_*,_1_, …, *x_r,nr_*} for *r* = 1, …, *R*. The state of each cell is located in a (measurable) space, *x* ∊ *ℳ*. We note that this space may not be the full dimension of the data set, but after a dimensionality reduction technique has been applied, e.g., PCA, diffusion maps etc. For example, in the case of RNA-Seq data, the state of a cell would consist of thousands of genes which would be too high to apply kernels to. We wish to find a probability distribution *ϱ* = *ϱ*(*t, x*) such that when *t* = *t_r_*, the probability of observing cells in states ***X**_r_* would be highly probable for *r* = 1, …, *R*.

Immediately necessary to ensure conservation of mass, we require 

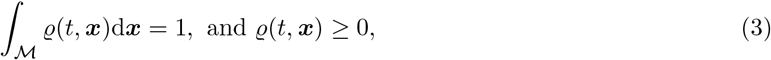

for all *t* ∊ (*t*_1_, *t_R_*). We now make the crucial assumption that our method relies on: each cell follows a (well behaved) autonomous dynamical system — implicit in this assumption is that cells do not interact, alternatively cell interactions can be accounted for via stochastic noise terms. Under these assumptions, we can interpret *ϱ*(*t, x*) Δ*x* as the probability a randomly selected cell has state in the interval [*x, x* + Δ*x*) at time *t*. We write down the (continuous-time) Perron–Frobenius equation for the dynamics of the density profile as 

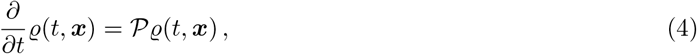

for initial condition *ϱ*(*t* = 0, *x*) = *ϱ*_0_(*x*). The 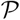 term is the continuous-time Perron–Frobenius operator [19, 20]. This operator is known by many names depending on the underlying dynamical system for the state evolution *x*(*t*) ∊ *ℳ* for *t* ∊ (*t*_1_, *t_R_*). For example, within SDEs equation (4) is a second order parabolic PDE known as the Fokker–Planck equation [28]; and for chemical reaction networks equation (4) is a system of coupled ODEs known as the chemical master equation [29].

### Finite Dimensional Approximation

We would like to find a finite dimensional approximation of of *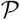*; we can do this with non-negative basis functions *ψ*(*x*) = [*ψ*_1_(*x*), … *ψ*,*_N_* (*x*)]^τ^. We take the ansatz that for all *t* ∊ (*t*_1_, *t_R_*), we can expand *ϱ* as the linear combination 

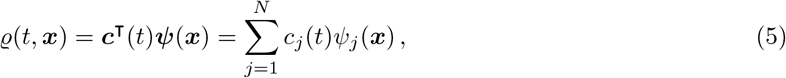

where *c*(*t*) = [*c*_1_(*t*), …, *c_N_* (*t*)]^τ^. To ensure the probability density integrates to one, we require *c* ∊ Λ where 

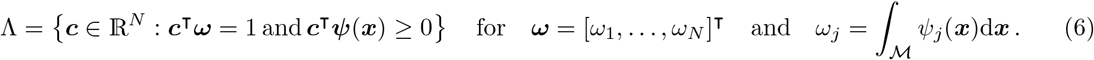

If the basis functions are themselves probability density functions, then Λ is the probability simplex. In Figure 1(a), we show how a distribution can be represented as a sum of normal distributions, that is, the density is given by a Gaussian mixture model.

We can derive a linear system of ODEs for the coefficients *c*(*t*). We do this by noting in weak form [30, 31] equation (4) is 

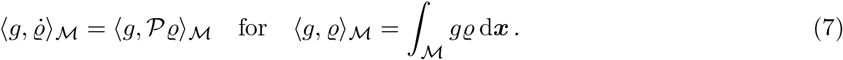

Choosing *g* = *ψ_i_* and expanding *ϱ* as in equation (5), then 

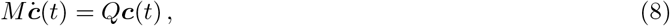

where *M_ij_* = 〈*ψ*_i_, *ψ*_j_〉 and 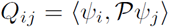. Assuming *M* is invertible, define 

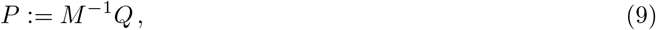

which is the projection of *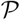* onto the basis functions {*ψ*_1_, …, *ψ_N_*}. That is, for *g*(*x*) = *c*^τ^ (*x*), we have the equality 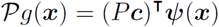. In order to preserve probability density and positivity, we require *P ∊ 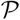* for 

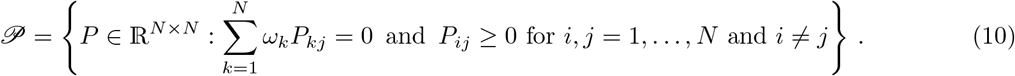

The explanation behind equation (10) is contained in the Appendix.

We can solve the dynamics for equation (4) using the approximation in equation (5) by using the matrix exponential operation, specifically 

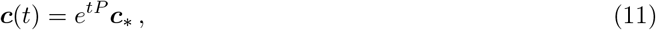

where *c_*_* are the coefficients at corresponding to the chosen initial condition 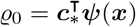.

### Selection of *P* matrix

We now need to address the issue of how to determine *P* from data. Consider a linear operator *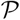* with matrix representation *P* on the space spanned by ψ. The *L*^2^ norm gives a measure of how well *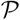* represents the evolution of the densities.

For an initial condition of 

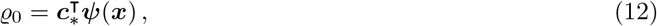

the squared relative prediction error at time *t* = *t_r_* for *r* = 1, …, *R* is 

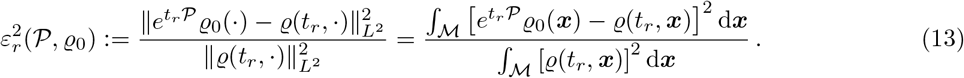

Notice that we specify that the error is a function of both the Perron–Frobenius operator and the initial condition, which is then treated as a parameter of the model. We then define the mean squared relative prediction error as 

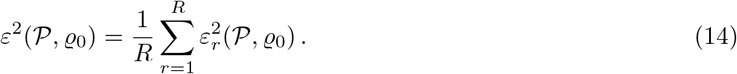

It would be ideal now to find 

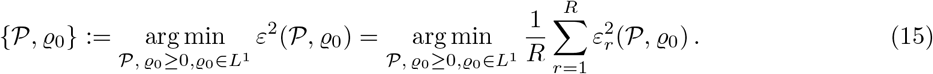

Of course, we do not know what this error is without using our finite dimensional approximation; therefore, by using equation (5) we calculate the time *t* = *t_r_* error as 

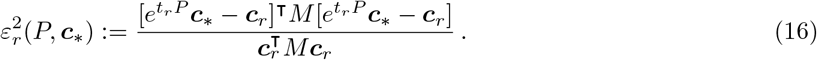

and therefore, our objective function is modified by using this finite dimensional approximation to 

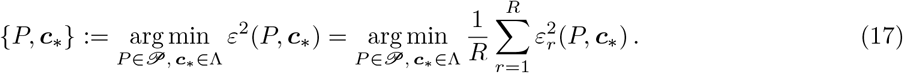

Unmentioned at this point is that: for a large quantity of basis functions the size over which this optimisation problem occurs is challenging. That is: one has *N*^2^ free parameters in the matrix *P*, with zero column sums one has *N*(*N* − 1) degrees of freedom; but one also has the initial condition to choose adding *N* parameters, so *N* − 1 degrees of freedom with unit column sum — in total (*N* − 1)(*N* + 1) degrees of freedom. Therefore, the problem is unapproachable without gradient calculations to speed up the optimization algorithm. Using the exponential matrix derivative (see Appendix), one can calculate the *t* = *t_r_* relative error with respect to *P* as 

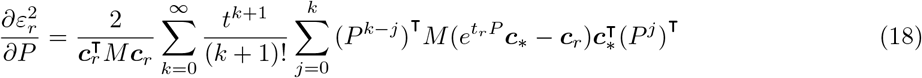

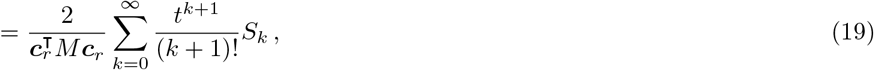

and the derivative with respect to *c_*_* as 

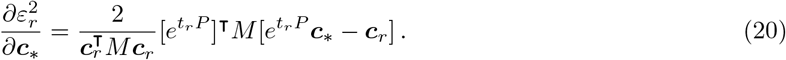

#### Algorithm 1 Algorithm to Determine Rate Matrix

**Require:** Data 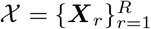 and observation times *t*_1_ < … < *t_R_*.

**Require:** Choose basis functions *ψ*(*x*) = [*ψ*_1_(*x*), …, *ψ_N_*(*x*)]^τ^.

^1:^ Solve 

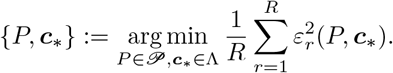

^2:^ **return** *P*

The terms 

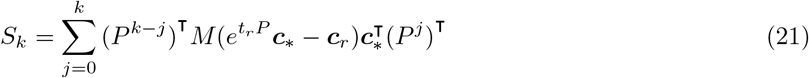

can be calculated using the recursion relation 

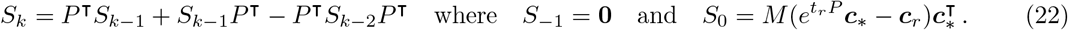

### Lasso Regularisation

For display purposes, we can promote sparsity in *P* by using a Lasso regularisation. We modify the error term in equation (17) to 

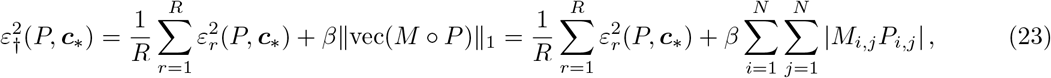

where º denotes the Hadamard product (or entrywise product) and vec(·) denotes the vectorisation of a matrix. In the case where the basis functions are probability density functions, we can calculate the derivative of this expression as 

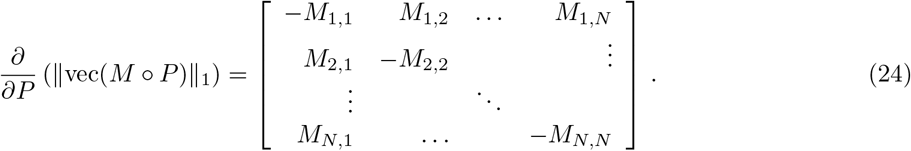

By using the mass matrix *M* as a weighting in front of the Perron–Frobenius matrix *P*, we are promoting edges between basis functions located apart from each other.

## Graph Visualisations

### *P* matrix visualisations

One can interpret the matrix *P* as a Markov rate matrix, in which case the entry *P_i,j_* shows the rate at which state *j* transitions into state *i*. Therefore, a cell in cluster *i* will switch to cluster *j* in time interval [*t, t* + Δ*t*) for Δ*t* > 0 with probability 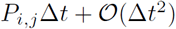. To reiterate, cells exist in states between the basis functions so instead of being at a single state, they are in a state which is a weighted combination of the basis functions. However, this interpretation allows us to plot a directed network with weighted adjacency matrix *P_i,j_*. Nodes can then be placed using one of a multitude of algorithms, in our case we use force-directed node placement with weights inversely proportional to the mass matrix *M*. We also plot the size of node *i* proportional to *P_i,i_* (as this is the rate at which state *i* remains in state *i*).

### Eigenfunction visualisation

To investigate key dynamical behaviours of a linear operator, a common theme is the study of the corresponding eigenproblem. By solving the eigenproblem, one can decompose the solution of the operator into components (known as eigenfunctions or eigenvectors) that will dynamically change with respect to the eigenvalue. By studying the eigenproblem, one can break down the solution into key behaviours and find important transitionary states.

For an eigenfunction satisfying 

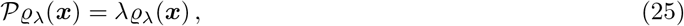

using the finite dimensional Galerkin approximation, there is the corresponding eigenvector 

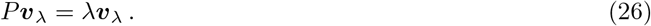

In low dimensions, one can simply plot this function as a linear combination 

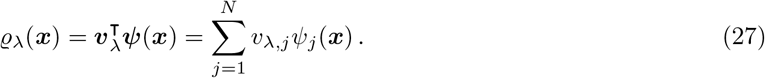

To ensure consistent scales when plotting, we demand 

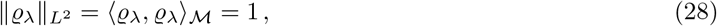

or in our finite dimensional representation 

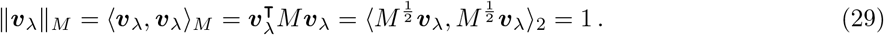

In high dimensions, our ability to visualise functions is limited. However, we have represented the function as a linear combination of basis functions and so we only need to present the coefficients in the eigenvector. To visualise if the eigenvector values have similar or dissimilar values, we can consider representing the eigenfunction as a graph. We specify the adjacency matrix for an undirected weighted graph as the outer product 

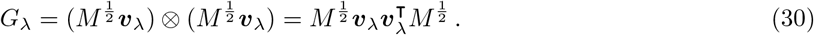

For Perron–Frobenius eigenfunctions with eigenvalue λ ≠ 0, the function will have both a positive part and a negative part (indicating where probability mass is flowing from and to). Therefore, by examining *G_λ_*, a positive value in entry (*i, j*) in *G_λ_* indicates basis functions *i* and *j* both have the same sign (positive or negative) and a negative value indicates they have opposite signs. By using the weighting of 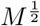 in front of the eigenvector *v_λ_* we ensure that the sizes of the eigenvectors are bounded. Occasionally it is the case that we get complex eigenvalues, in which case they appear as complex conjugates and one can plot the real and imaginary parts separately.

## Conclusion

We presented Dynamic Distribution Decomposition for identification of dynamically important states of biological processes. This method operates on snapshot time series data and infers dynamically important states by mapping between distributions. We applied our approach to synthetic data generated for a simple test system, and then further to a mass cytometry time series data set for iPSC reprogramming. Our approach performed well for both systems showing key dynamical states. For the experimental system of iPSC reprogramming of fibroblasts, we could also identify key time points where the current experimental set-up is insufficient elucidate the reprogramming process and where further investigation is warranted.

DDD can be computed efficiently, e.g., via the recursion relation given by equation (55), but we have not optimised the implementation. In the case where there is more than 100 basis functions, our minimisation procedure using the inbuilt MATLAB multivariate minimisation algorithm can be unreasonably slow; therefore, we will investigate more efficient implementations.

DDD depends on a few design decisions, such as the choice of basis functions. In this manuscript, we used basis functions as components of a Gaussian mixture model and gave parameters that needed tuning to alter the fit. Our method for choosing basis functions does not have an optimal configuration with regards to minimising error. This is because by sending the regularisation value to infinity one obtains perfect fits, and sending the regularisation value to zero can lead to ill fitting solution (basis functions with same mean etc). However, one can use the parameter as a rule of thumb to ensure enough basis functions are included. For other applications other choices in basis functions are conceivable, for example, radial basis functions [32]; piecewise linear basis functions [33]; and global basis functions [19, 25] to name a few. When the basis functions have finite support, the mass matrix *M* will be sparse, in which case the Lasso step will not be necessary. It would also make sense to use basis functions built around the data type, for example negative binomial distributions are often used to model UMI counts from single-cell RNA-Seq data. When one uses a single basis function centred around each data point, one refers to this as kernel density estimation, of which there are optimal methods to choose the basis function [34]; when using a small number of basis functions for a large number of sample points, there are likely optimal ways of choosing them which we will investigate in future work.

DDD could be applied to investigate pseudo-time ordered single-cell data of single time point experiments. Here, one uses single-cell data measured at only a single time point to carry out trajectory inference and subpopulation identification to infer biological processes, e.g., the cell cycle; a review of such methods can be found in Ref. [35]. It may be possible to improve our fits by combining approaches: while cells are monitored with regards to experimental time, individual cell time coordinates might deviate due to asychronity of process initiation; this could be incorporated to get smoother Perron–Frobenius operators between time points, see Ref. [36]. This would then be a biologically motivated method to account for delays in the system.

The work presents Dynamic Distribution Decomposition, linking operator theory to the practical world of high-dimensional data analysis. While we focus on application of DDD to mass cytometry measurements, it is conceivable to expand to applications to single-cell RNA sequencing time series as well as biological processes other than an iPSC reprogramming. We expect DDD and method variations will be instrumental in providing intuitive understanding of such biological processes.

## Acknowledgments

We gratefully acknowledge the support of Anna Klimovskaia, Will Macnair, Ioana Sandu, Dario Cerletti, and Eli Zunder. J.P.T-K was supported by the medical research and development HDL-X grant from SystemsX.ch. A.N.R. is partially supported by the EPSRC Centre For Doctoral Training in Industrially Focused Mathematical Modelling (EP/L015803/1).

## Additional Mathematical Details

### Choice in basis functions

One option for choosing the basis functions is to use probability density functions, so 

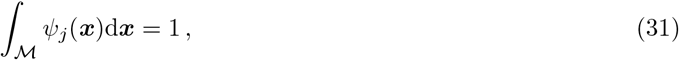

for *j* = 1, …, *N*. We can find the values of *c* at the observed time points by noting that the value of the coefficient at time *t* = *t_r_* must be proportional to the probability that basis function *j* created the data at that time point, so 

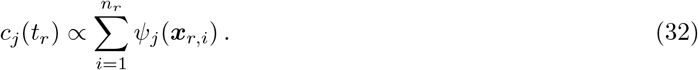

One can then normalise 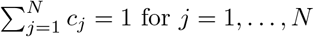 to find the coeffient vector *c_r_*. Assuming multivariate Normal distributions as basis functions, we have 

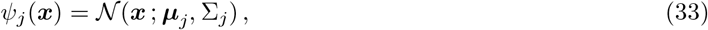

where 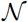 is the probability density function for a multivariate Normal distribution with mean *µ* and covariance matrix ∑. To determine the mass matrix, adapting results from Ref. [37] we can analytically calculate the inner product between two multivariate normal distributions as 

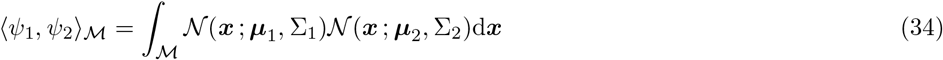

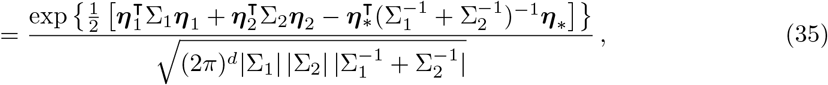

for 

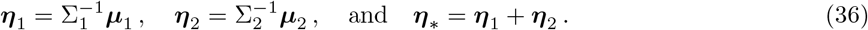

### Requirements on Perron–Frobenius matrix approximation

To preserve probability, it must be the case that 

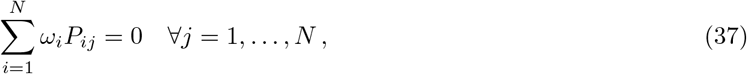

which is identical to the condition for continuous time Markov chains. This is because we require 

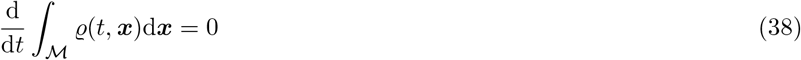

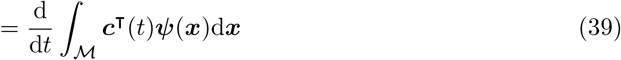

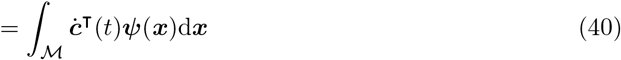

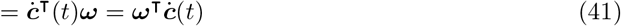

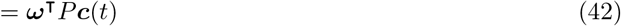

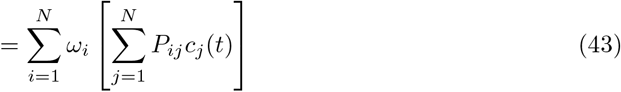

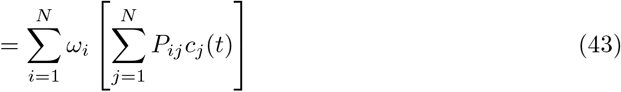

Therefore, for mass to be conserved, equation (10) holds.

We also require the operator 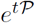 preserves positivity, that is, 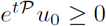 for functions *u*_0_ ≥ 0. This implies that the off-diagonal entries of *P* must be non-negative. The reason is as follows. As *ψ* ≥ 0, the function 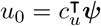 must have *c_u_* ≥ 0 due to linear independence of the normalised basis functions. It therefore follows that *e^tP^* ≥ 0 element wise, because the positivity condition and linear independence of the normalised basis functions require that 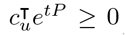 element wise. Thus, 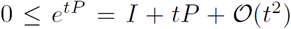 for all *t* ≥ 0, which in particular means that 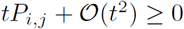 for *i* ≠ *j*. The condition *P_i,j_* ≥ 0 follows by considering arbitrarily small *t* > 0. This is sufficient for *e^tP^* to be element wise non-negative, as the matrix exponential of a matrix with non-negative off-diagonal elements is always non-negative element wise. Note that this, together with the mass conservation condition, implies that the diagonal entries of *P* are all negative.

### Gradient Calculation

For relevant background reading on functions of matrices, see Ref. [38]. Let *ℋ* denote the Hilbert space ℝ^*N*^ with inner product 〈***p, q***〉_*ℋ*_ = *p*^τ^ *M **q**, P* ∊ ℝ^*N×N*^. For *t* > 0, and ***p, q*** ∊ *ℋ*, we derive the gradient of 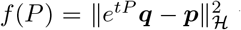 that is used for the gradient calculation in Equation (18). Let *D_f_* (*P*; *dP*) be the directional derivative of *f* at *P*, in the direction *dP*. Let 〈·, ·〉*_F_* denote the Frobenius inner product on ℝ^*N×N*^. The gradient of *f* is the matrix *G* ∊ ℝ^*N×N*^ such that *D_f_*(*P*; *dP*) = 〈*G, dP*〉_F_ for any *dP* ∊ ℝ^*N×N*^.

First, note that 

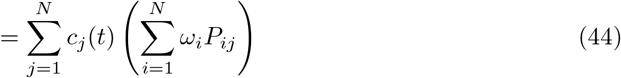

We therefore need an expression for the directional derivative *D_e^t·^_* (*P*; *dP*) of *P* → *e^tP^*. By definition of the matrix exponential, we have that 

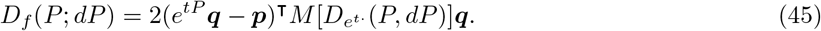

The directional derivative is thus 

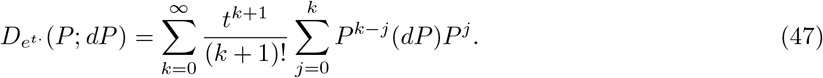

By combining (45) and (47) we see that the directional derivative of *f* is 

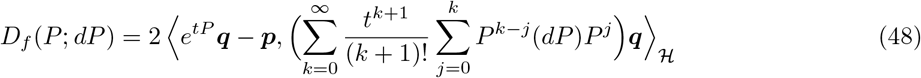

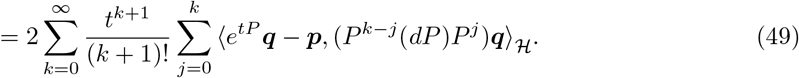

The final step is to derive *G* such that *D_f_*(*P*; *dP*) = 〈*G, dP*〉*_F_*. Note that 〈***q**, A**p***〉_*ℋ*_ = 〈*M**qp***^τ^, *A*〉_*F*_ for arbitrary ***p, q*** ∊ ℝ^*N*^ and *A* ∊ ℝ^*N × N*^. Thus, (49) becomes 

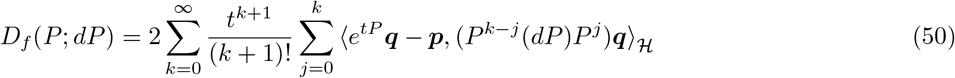

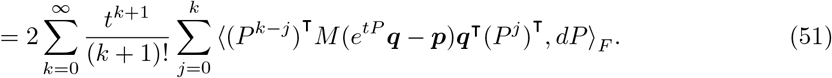

It therefore follows that the gradient of *f* at *P* is the matrix 

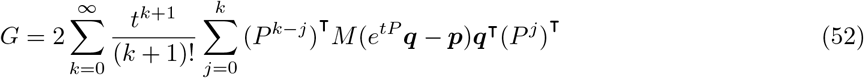

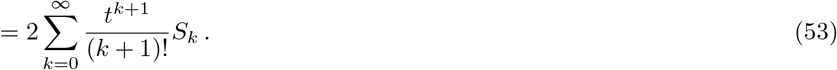

The terms 

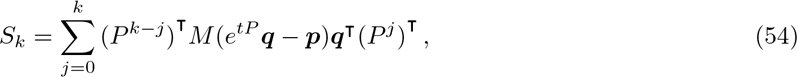

can be calculated using the recursion relation 

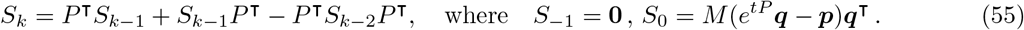

## Additional Figures

**Fig 8.**
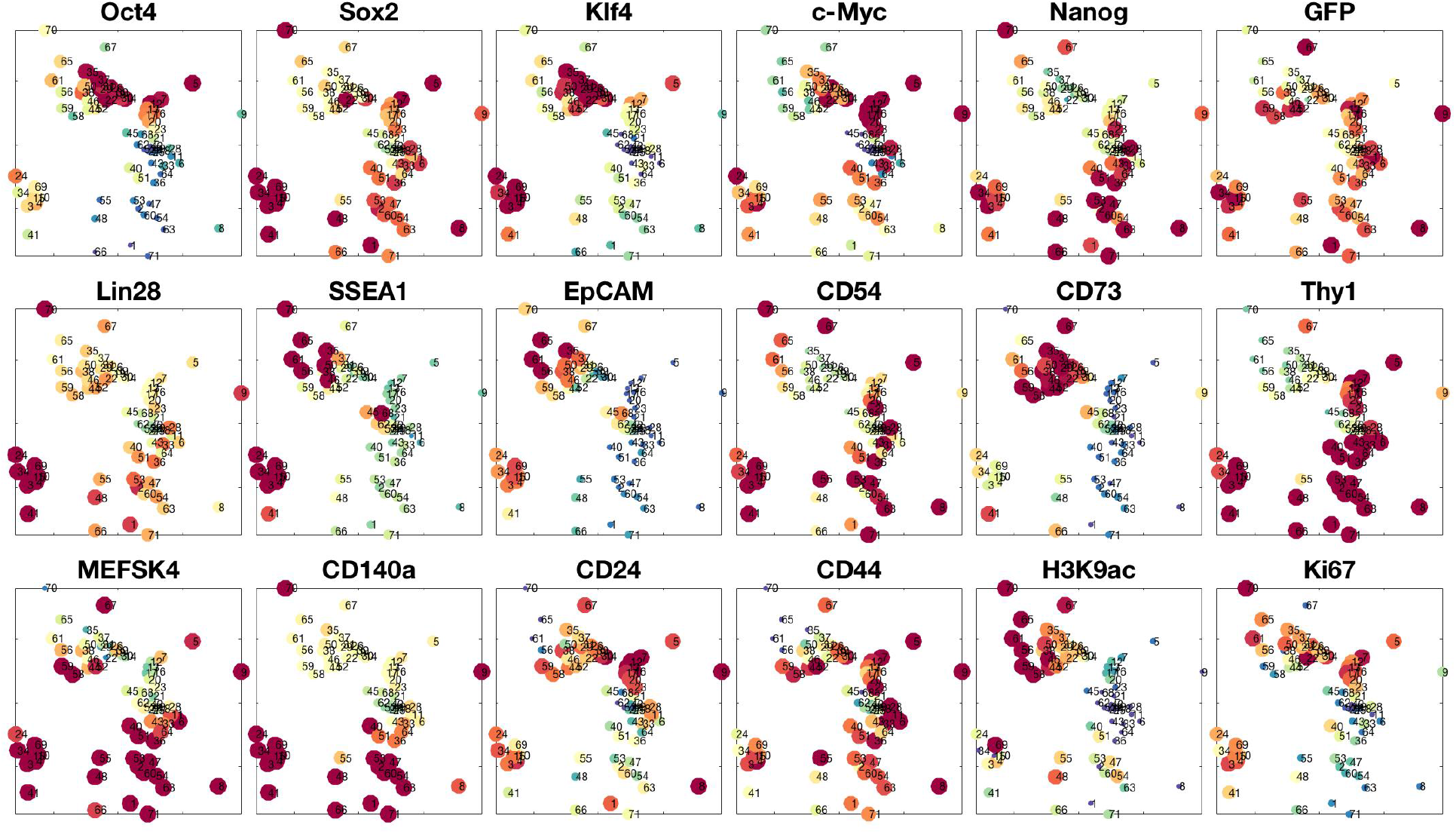
Protein expression of Nanog-Neo cell line data taken from Zunder *et. al*. [26]. Locations are plotted using positions of nodes in Figure 6. After the raw data is transformed via *f*(*x*) = arcsin(*x*/5) and *z*-scored, we limit the colour scale to the middle 50% of the data (lower quartile in dark blue, upper quartile in dark red).

## Footnotes

1 This numerical scheme coincides with a Milstein scheme (and is of order 1).

## References

1. Ziegenhain C, Vieth B, Parekh S, Reinius B, Guillaumet-Adkins A, Smets M, et al. Comparative analysis of single-cell RNA sequencing methods. Molecular cell. 2017;65(4):631–643.

2. Spitzer MH, Nolan GP. Mass cytometry: single cells, many features. Cell. 2016;165(4):780–791.

3. Zechner C, Ruess J, Krenn P, Pelet S, Peter M, Lygeros J, et al. Moment-based inference predicts bimodality in transient gene expression. Proceedings of the National Academy of Sciences. 2012;109(21):8340–8345.

4. Klimovskaia A, Ganscha S, Claassen M. Sparse regression based structure learning of stochastic reaction networks from single cell snapshot time series. PLoS computational biology. 2016;12(12):e1005234.

5. Pantazis Y, Tsamardinos I. A Unified Approach for Sparse Dynamical System Inference from Temporal Measurements. arXiv preprint arXiv:171000718. 2017;.

6. Schiebinger G, Shu J, Tabaka M, Cleary B, Subramanian V, Solomon A, et al. Reconstruction of developmental landscapes by optimal-transport analysis of single-cell gene expression sheds light on cellular reprogramming. bioRxiv. 2017;doi:10.1101/191056.

7. Ulam S. A collection of Mathematical Problems. vol. 8 of Interscience tracts in pure and applied mathematics. Interscience Publishers Inc.; 1960.

8. Lasota A, Mackey MC. Probabilistic properties of deterministic systems. Cambridge University Press; 1985.

9. Fokker AD. Die mittlere Energie rotierender elektrischer Dipole im Strahlungsfeld. Annalen der Physik. 1914;348(5):810–820. doi:10.1002/andp.19143480507.

10. Planck M. Über einen Satz der statistischen Dynamik und seine Erweiterung in der Quantentheorie. Reimer; 1917.

11. Koopman BO. Hamiltonian systems and transformation in Hilbert space. Proceedings of the National Academy of Sciences. 1931;17(5):315–318. doi:10.1073/pnas.17.5.315.

12. von Neumann J. Zur Operatorenmethode in der klassischen Mechanik. Annals of Mathematics. 1932; p. 587–642. doi:10.2307/1968537.

13. Neumann JV. Zusatze Zur Arbeit, Zur Operatorenmethode… Annals of Mathematics. 1932; p. 789–791. doi:10.2307/1968225.

14. Bachelier L. Théorie des probabilités continues. Journal de Mathématiques Pures et Appliquées. 1906;2:259–328.

15. Kolmogoroff A. Über die analytischen Methoden in der Wahrscheinlichkeitsrechnung. Mathematische Annalen. 1931;104(1):415–458. doi:10.1007/BF01457949.

16. Schmid PJ. Dynamic mode decomposition of numerical and experimental data. Journal of fluid mechanics. 2010;656:5–28.

17. Tu JH, Rowley CW, Luchtenburg DM, Brunton SL, Kutz JN. On Dynamic Mode Decomposition: Theory and applications. Journal of Computational Dynamics. 2014;1(2):391–421.

18. Williams MO, Kevrekidis IG, Rowley CW. A data–driven approximation of the Koopman operator: Extending dynamic mode decomposition. Journal of Nonlinear Science. 2015;25(6):1307–1346.

19. Klus S, Koltai P, Schütte C. On the numerical approximation of the Perron-Frobenius and Koopman operator. Journal of Computational Dynamics. 2016;3(1):51–79. doi:10.3934/jcd.2016003.

20. Klus S, Nüske F, Koltai P, Wu H, Kevrekidis I, Schütte C, et al. Data-driven model reduction and transfer operator approximation. arXiv preprint arXiv:170310112. 2017;.

21. Proctor JL, Eckhoff PA. Discovering dynamic patterns from infectious disease data using dynamic mode decomposition. International health. 2015;7(2):139–145.

22. Proctor JL, Brunton SL, Kutz JN. Dynamic mode decomposition with control. SIAM Journal on Applied Dynamical Systems. 2016;15(1):142–161.

23. Brunton BW, Johnson LA, Ojemann JG, Kutz JN. Extracting spatial–temporal coherent patterns in large-scale neural recordings using dynamic mode decomposition. Journal of neuroscience methods. 2016;258:1–15.

24. Mauroy A, Goncalves J. Koopman-based lifting techniques for nonlinear systems identification. arXiv e-prints. 2017;.

25. Riseth AN, Taylor-King JP. Operator Fitting for Parameter Estimation of Stochastic Differential Equations. arXiv e-prints. 2017;.

26. Zunder ER, Lujan E, Goltsev Y, Wernig M, Nolan GP. A continuous molecular roadmap to iPSC reprogramming through progression analysis of single-cell mass cytometry. Cell Stem Cell. 2015;16(3):323–337.

27. Akaike H. A new look at the statistical model identification. IEEE transactions on automatic control. 1974;19(6):716–723.

28. Øksendal B. Stochastic Differential Equations: An introduction with applications. 5th ed. Springer Verlag; 2000.

29. Erban R, Chapman J, Maini P. A practical guide to stochastic simulations of reaction-diffusion processes. arXiv preprint arXiv:07041908. 2007;.

30. Ziemer WP. Weakly differentiable functions: Sobolev spaces and functions of bounded variation. vol. 120. Springer Science & Business Media; 2012.

31. Gilbarg D, Trudinger NS. Elliptic partial differential equations of second order. Springer; 2015.

32. Fornberg B, Flyer N. A primer on radial basis functions with applications to the geosciences. SIAM; 2015.

33. Alberty J, Carstensen C, Funken SA. Remarks around 50 lines of Matlab: short finite element implementation. Numerical Algorithms. 1999;20(2–3):117–137.

34. Botev ZI, Grotowski JF, Kroese DP, et al. Kernel density estimation via diffusion. The annals of Statistics. 2010;38(5):2916–2957.

35. Saelens W, Cannoodt R, Todorov H, Saeys Y. A comparison of single-cell trajectory inference methods: towards more accurate and robust tools. bioRxiv. 2018;doi:10.1101/276907.

36. Fischer DS, Fiedler AK, Kernfeld E, Genga RMJ, Hasenauer J, Maehr R, et al. Beyond pseudotime: Following T-cell maturation in single-cell RNAseq time series. bioRxiv. 2017;doi:10.1101/219188.

37. Bromiley P. Products and convolutions of Gaussian probability density functions. Tina-Vision Memo. 2003;3(4):1.

38. Higham NJ. Functions of matrices: theory and computation. vol. 104. Siam; 2008.

